# Significant improvement of environmental DNA assay by targeting retrotransposon sequences characteristic to *Anguilla* eels

**DOI:** 10.1101/2025.03.06.641759

**Authors:** Itsuki T. Hirayama, Yuta Kunimasa, Aya Takeuchi, Toshifumi Minamoto

**Affiliations:** Graduate School of Human Development and Environment, Kobe University; Japan Society for the Promotion of Science; 3Department of Fisheries, Faculty of Agriculture, Kindai University, 3327-204 Nakamachi, Nara 631-8505, Japan

**Keywords:** *Anguilla*, Environmental DNA, Retroelements, Short interspersed nucleotide elements (SINE)

## Abstract

Environmental DNA (eDNA) analysis faces challenges regarding sensitivity and quantification accuracy, particularly in areas with low target species densities or during seasons when eDNA release decreases. Multi-copy markers such as mitochondrial genomic DNA and ribosomal DNA (rDNA) have been widely used in eDNA analysis to address this issue. However, the copy number of the DNA markers per cell remains a potential bottleneck for eDNA sensitivity. In this study, we aimed to increase the sensitivity of eDNA assays by using retrotransposons, which are abundant in the genome, as novel target markers. We developed an assay targeting UnaSINE1, a short interspersed nucleotide element (SINE) characteristic of *Anguilla* eels, and compared its sensitivity and accuracy with that of an established mitochondrial 16S rRNA marker. Our results demonstrated that UnaSINE1 was detected at over 100 times the copy number of the mitochondrial marker in both genomic and eDNA samples. In the river surveys, the 16S marker was positive in 32 of the 81 samples, whereas the UnaSINE1 marker was positive in 62 samples, indicating that the use of SINE marker remarkably reduced false negatives. Furthermore, both biological and technical replicates exhibited improved positive consistency and reduced variability in quantification, leading to more robust presence/absence determination and quantitative results. Utilizing retrotransposon sequences as markers requires additional effort for sequence acquisition and organization and may limit taxonomic resolution to the genus level. However, this approach significantly improves sensitivity without increasing the labor or cost of sampling and PCR analysis, making it highly practical for eDNA studies.

## 1. Introduction

Rapid loss of biodiversity has increased the importance of biological monitoring. Traditionally, biological monitoring has been conducted primarily through physical captures and visual surveys. Environmental DNA (eDNA) analysis, which examines DNA fragments that remain in the environment, has gained widespread use in recent years. This method, which involves collecting one to several liters of water from rivers or oceans and analyzing the DNA within, offers several advantages. It minimizes the labor required at survey sites, is non-invasive and non-destructive, and enables detecting a wide range of species over large areas. However, due to insufficient detection sensitivity, eDNA analysis has limitations, particularly the issue of false negatives. False negatives can occur when the population density is extremely low or, the target species releases only small amounts of eDNA (Millard-Martin et al., 2024; Schultz et al., 2015). Additionally, eDNA detection is challenging in water bodies with high levels of inhibitory substances, and in winter when biological activity decreases (Buxton et al., 2018; Hunter et al., 2019; McKee et al., 2015). Several approaches have been attempted in standard eDNA analysis, including increasing the water sample volume, improving DNA concentration and purification, and using high-sensitivity PCR platforms such as digital PCR, to increase detection sensitivity (Capo et al., 2021; Schabacker et al., 2020; Williams et al., 2017). However, these additional measures increase labor and costs, limiting their practical application (Hinlo et al., 2017; Sanches et al., 2020). Moreover, when the actual concentration of eDNA in a given environment is extremely low, there is a risk that no target molecular markers will be present in the collected sample, rendering subsequent processing ineffective. Therefore, the use of multi-copy markers is essential to improve the sensitivity of eDNA detection. To date, mitochondrial genome DNA and tandemly repeated ribosomal RNA (rRNA) genes on chromosomes (rDNA) have been widely used as eDNA markers (Shu et al., 2020; Minamoto et al., 2017; Jo et al., 2021). However, the number of copies of these markers in a single cell remains a limiting factor for eDNA sensitivity (Xia et al., 2021).

Therefore, we focused on retrotransposons, which are present at much higher copy numbers within cells than mitochondrial or rDNA markers, to overcome this limitation. Retrotransposons are mobile genetic elements that propagate by “copying and pasting” sequences throughout a genome (Kramerov & Vassetzky, 2005). They play a crucial role in the evolution and complexity of eukaryotic genomes (Cordaux & Batzer, 2009; Almojil et al., 2021). Notably, their copy numbers are extraordinarily high. The human genome contains approximately 850,000 copies of long interspersed elements (LINEs) and 1.5 million copies of short interspersed elements (SINEs), which collectively account for approximately 34% of the genome (Lander et al., 2001). This suggested that using retrotransposons as eDNA markers could improve detection sensitivity by more than 100-fold compared with using conventional markers. Previous studies on fish have identified species-specific retrotransposons, such as UnaSINE1 in *Anguilla* eels and SmaI in salmon, which have been used for phylogenetic analyses across species (Kajikawa et al., 2004; Matveev et al., 2009).

Eels of the genus *Anguilla* are catadromous fishes that spawn in the ocean and migrate to rivers and coastal areas in temperate and tropical regions for growth (Arai, 2020). Three *Anguilla* species are known from Japan: *Anguilla japonica* (Japanese eel), *Anguilla marmorata*, and *Anguilla bicolor pacifica*. Among these, *A. japonica* is widely distributed, except in the subarctic regions, whereas the tropical species *A. marmorata* and *A. bicolor pacifica* are primarily found in southern Japan (Tesch, 2003). *Anguilla japonica* has long been used as a food resource in East Asia, but its population decline has raised concerns (Kuroki et al., 2014; Kaifu & Yokouchi, 2019). Consequently, *A. japonica* has been listed as an endangered species (EN) on the IUCN Red List (IUCN, 2020).

The accurate assessment of *Anguilla* populations requires detailed monitoring, and eDNA analysis is a promising method for surveying this widely distributed, low-density species (Kasai et al., 2021). Previous studies have shown that eDNA analysis can detect eels more efficiently and cost-effectively than traditional capture surveys in river environments, allowing biomass estimation (Itakura et al., 2019; 2020a). However, in a study conducted in the spawning area of *A. japonica* in the southern part of the West Mariana Ridge, only 3 out of 117 samples tested positive, and eDNA was detected in only one of three PCR replicates for each sample (Takeuchi et al., 2019), indicating the limitations of standard eDNA methods.

Therefore, we developed a novel assay targeting UnaSINE1, a known *Anguilla*-specific SINE, to address these limitations. We applied this assay to water samples from rivers in the Hyogo Prefecture, Japan, where *A. japonica* is present, and compared its performance with that of the commonly used 16S mitochondrial marker.

## 2. Materials and methods

### 2.1 Assay Development

The sequence information for UnaSINE1 was retrieved from the NCBI database (https://www.ncbi.nlm.nih.gov/). We used UnaSINE1-8 (Accession: AB179628.1) as a representative query and conducted a BLAST search against all nucleotide sequences (nr/nt) and genome assemblies (refseq_genomes) available for the order Anguilliformes. All identified sequences were collected, and sequences shorter than 200 bp were excluded before sequence alignment was performed. Primers targeting conserved core sequences specific to *Anguilla* were designed manually based on the alignment data. The primer design criteria were as follows: amplicon size of 80–200 bp, primer length of 15–30 bp, melting temperature (Tm) around 60 °C, a Tm difference between forward and reverse primers of ≤2 °C, and GC content of 30–70%.

### 2.2 Specificity Test of Assay Using Genomic DNA (gDNA)

We first performed an *in silico* specificity test using Primer-BLAST with default settings to verify that the designed assay did not amplify DNA from non-Anguilla species. Subsequently, an *in vitro* specificity test was conducted using the available genomic DNA (gDNA) from two *Anguilla* species inhabiting Japan (*A. japonica* and *A. marmorata*) and another marine eel species (*Conger myriaster*). gDNA was extracted from the tissue samples using the DNeasy Blood & Tissue Kit (Qiagen, Hilden, Germany). The extracted gDNA was diluted to a final concentration of 5 pg/µL (equivalent to 10 pg/reaction) in TE buffer (pH 8.0) and used as a PCR template.

Real-time PCR was performed using the StepOnePlus Real-Time PCR System (Thermo Fisher Scientific, Waltham, MA, USA). Triplicate reactions (20 µL each) were conducted for each sample. The reaction mix contained 1x TaqMan Environmental Master Mix 2.0 (Thermo Fisher Scientific), 0.1□μL of AmpErase Uracil N-glycosylase (Thermo Fisher Scientific), 900□nM of each primer, 125□nM of probe, and 2 μL template DNA. The thermocycling profile was 50 °C for 2□min and 96 °C for 10□s, followed by 55□cycles of 96 °C for 15□s and 60 °C for 1□min. The primers and probes used were either the mitochondrial 16S marker (Watanabe et al. 2004) or a newly developed SINE marker (Table 1). A negative control was included for each PCR run to confirm the absence of cross-contamination.

**Table 1.**
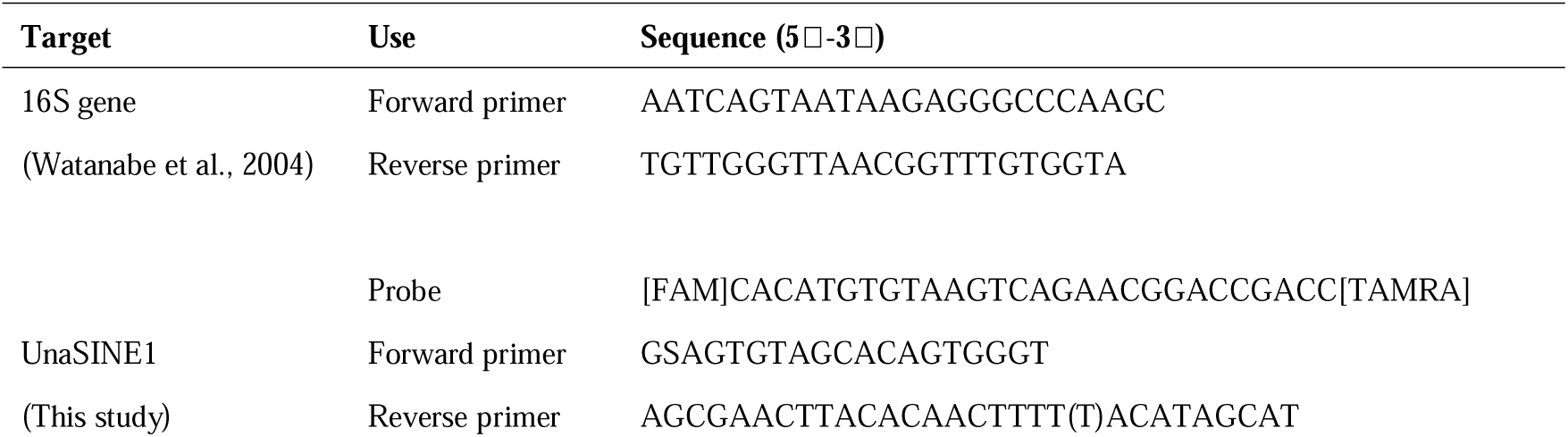
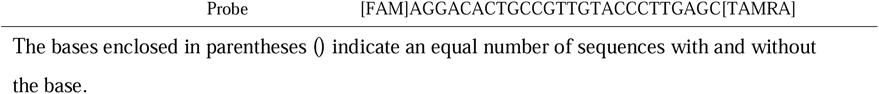
The sequences of the primers and probes.

### 2.3 Sensibility Test

Serial dilutions of *A. japonica* gDNA at concentrations ranging from 10□^4^ to 10^1^ pg/reaction were prepared to compare the limits of detection (LOD) and quantification (LOQ) of the mitochondrial and SINE markers. The same PCR conditions as those used for the specificity test (Section 2.2) were used with three negative controls included in each run. The LOD was defined as the lowest concentration at which at least one of the three replicates tested positive, while the LOQ was defined as the lowest concentration at which all three replicates tested positive (Jo et al., 2024).

### 2.4 Field Survey

Water sampling was conducted at nine freshwater sites in each of the rivers of the Hyogo Prefecture, Japan, in February, June, and October 2023, yielding a total of 27 points (Fig. 1). At each sampling point, three 500 mL water bottle samples were collected as biological replicates. A negative field control was used in each survey to confirm the absence of cross-contamination. Benzalkonium chloride (BAC, 0.1% w/v) was added to the water samples and the control before being transported to the laboratory to prevent DNA degradation (Yamanaka et al., 2017). Samples were filtered using GF/F filters (GE Healthcare Life Science, Little Chalfont, UK) and stored at −28 °C. DNA was extracted from the filters using a DNeasy Blood & Tissue Kit (Qiagen) following the protocol outlined in the eDNA Society’s experimental manual (Minamoto et al., 2021). A total of 100 µL of eDNA solution was obtained. Real-time PCR was performed on the extracted eDNA using assays for both the SINE and mitochondrial 16S markers. The PCR conditions were identical to those used in the specificity test (Section 2.2), and the reactions were conducted using the QuantStudio 3.0 Real-Time PCR System (Thermo Fisher Scientific). Synthetic DNA fragments (Eurofins Genomics, Tokyo, Japan) of the consensus sequence of *A. japonica* UnaSINE1 (Table S1) were used as calibration standards at concentrations of 30, 300, 3,000, and 30,000 copies/reaction to quantify eDNA copy numbers. A bottle sample was considered positive if at least one of the three PCR (technical replicates) tested positive. A sampling point was considered positive if at least one of three bottle samples (biological replicates) tested positive. Non-detected replicates were assigned a value of 0 copies, and the mean eDNA concentration and coefficient of variation (CV) were calculated for both technical and biological replicates.

**Figure 1.**
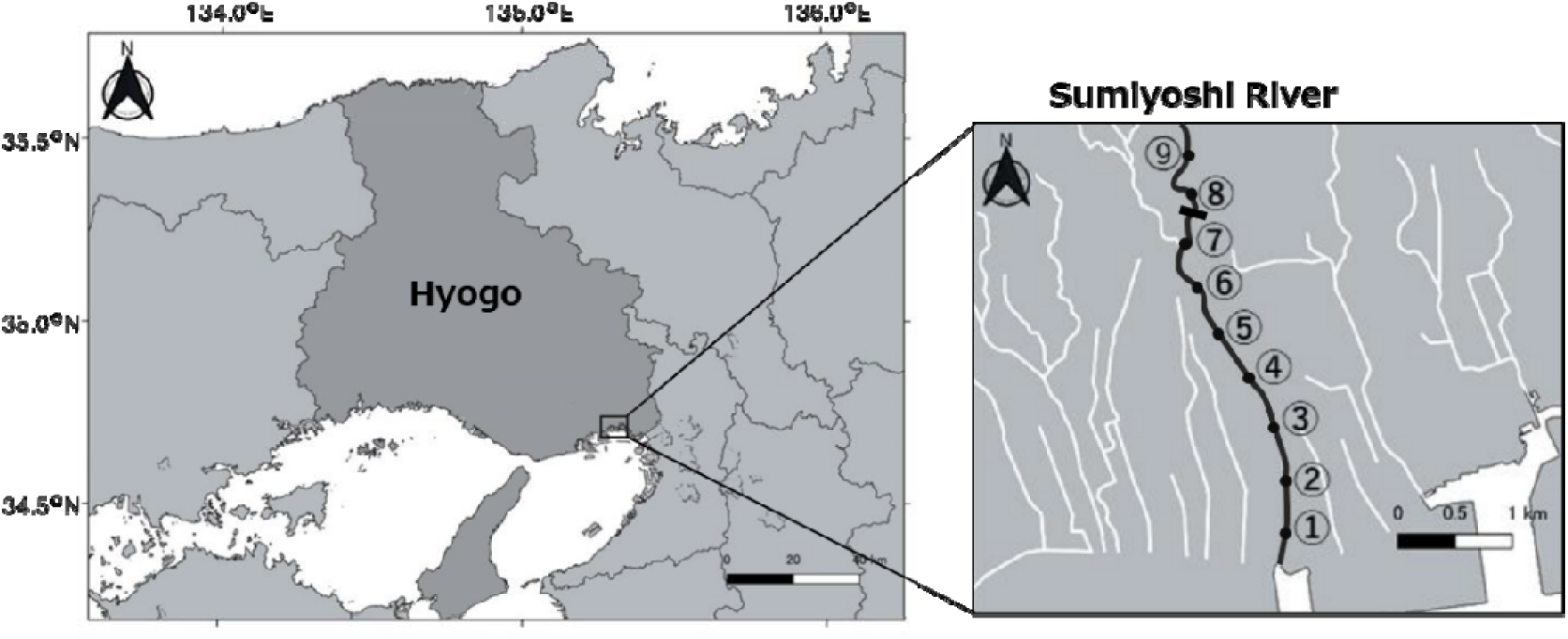
Geographic location of the Sumiyoshi River in southern Hyogo Prefecture, Japan, and the nine sampling sites where 500 mL water was collected. The black rectangle between Sites 7 and 8 indicates a weir more than 3 meters high.

### 2.5 Statistical Analysis

All statistical analyses were performed using R version 4.4.2 (R Core Team, 2024). The eDNA concentration per reaction (copies/2 µL template) was converted to the concentration per liter of water (copies/L). The eDNA concentration for each marker was log10-transformed (+1) to meet normality and analyzed using linear mixed models (LMMs). The explanatory variable was the sampling point, whereas the random effects included the sampling site and month (Table S1). Model II regression analysis was conducted to test the correlation between eDNA concentrations of the two markers. Sites 8 and 9, where no eDNA was detected during the study period, were excluded from the analysis.

The detection frequencies of the two markers were compared using McNemar’s test to evaluate whether the use of the SINE marker increased the detection probability at the sampling points. We examined the consistency of technical replicates among bottle samples in which the mitochondrial marker was detected using McNemar’s test to assess whether the SINE marker reduced detection errors. Additionally, Wilcoxon signed-rank tests were conducted to compare the CV values between SINE and mitochondrial markers for both biological and technical replicates.

Finally, differences in eDNA concentrations among sampling sites and months were analyzed separately for each marker using linear models (LMs), with the sampling point as the explanatory variable. Multiple comparisons were performed using post-hoc Tukey’s adjustments. Sites 8 and 9 were excluded from the analysis.

## 3. Results

We developed an assay targeting the UnaSINE1 cluster that is specific to the genus *Anguilla* (Table 1). The forward and reverse primers included degenerate sites or gaps to maximize the number of matched UnaSINE1 sequences. The assay amplified sequences from all *Anguilla* species with genome assemblies registered in the NCBI database in the *in silico* specificity test, whereas no amplification was observed in non-*Anguilla* species. Primer-BLAST results indicated that the amplicon length varied between 170 and 207 bp in *A. japonica*, with a median length of 185 bp. Real-time PCR successfully amplified DNA from both *A. japonica* and *A. marmorata* but not from *C. myriaster*, in the *in vitro* specificity test, confirming the specificity of the assay. When 10 pg of gDNA was used as a template, the difference in Ct values between *A. japonica* and *A. marmorata* was only 0.046, indicating a similar amplification efficiency for the two species.

Despite having a longer amplicon than the mitochondrial 16S marker (107 bp), the SINE marker exhibited significantly higher copy numbers in the template DNA. The LOD and LOQ of the SINE were two orders of magnitude lower than those of the mitochondrial 16S marker (Fig. 2). The LOQs were 0.01 pg/reaction and 1 pg/reaction for the SINE and 16S markers, respectively. The LODs were 0.001 pg/reaction and 0.1 pg/reaction for the SINE and 16S markers, respectively.

**Figure 2.**
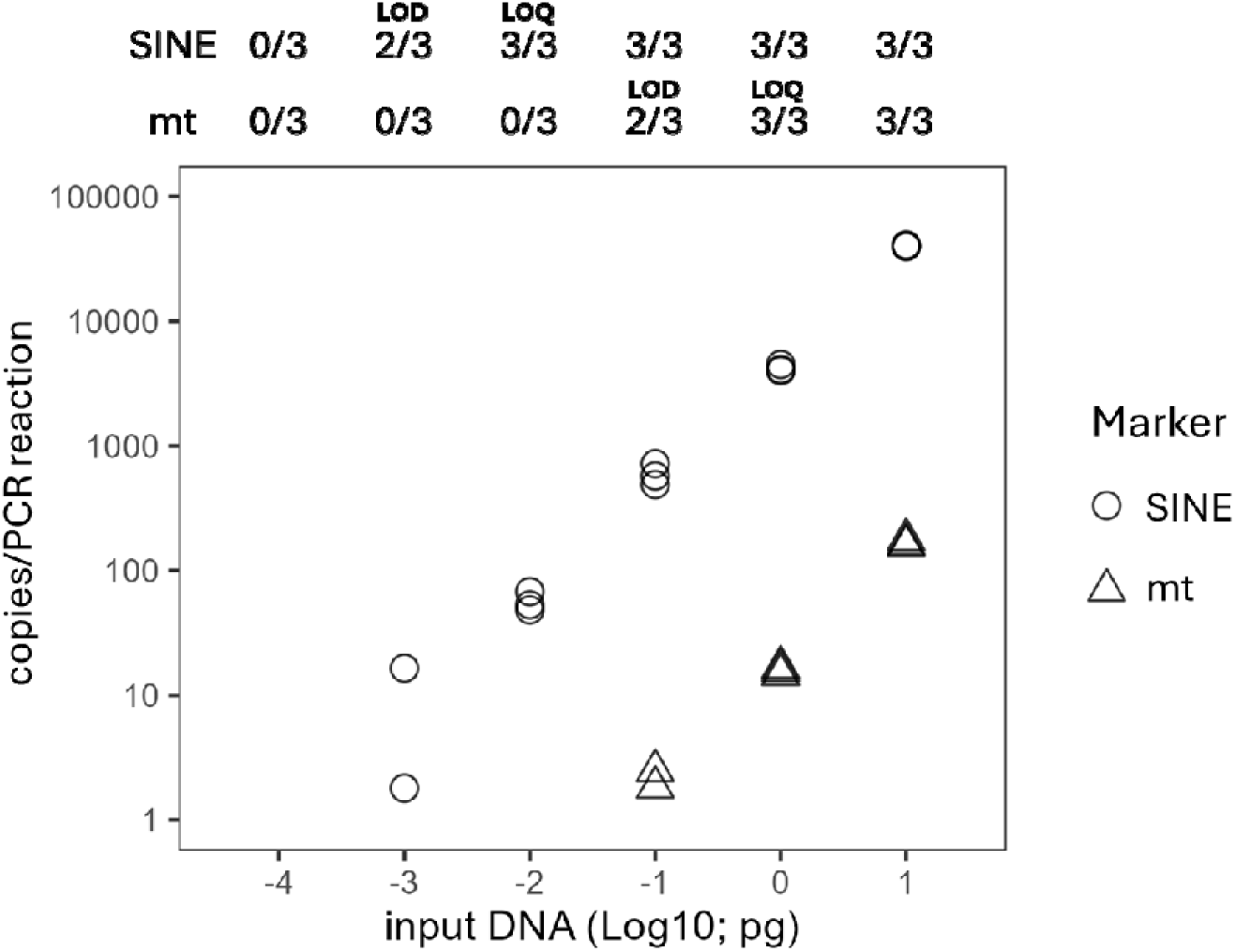
Results of the sensitivity test for the SINE (UnaSINE1) and mitochondrial 16S (mt) assays. The top panel shows the number of positive PCR replicates out of three for each concentration of genomic DNA. The y-axis represents the number of marker copies per PCR template (2 µL), calculated using the standard DNA calibration curve. The limit of quantification (LOQ) was defined as the lowest input DNA concentration at which all three replicates tested positive, while the limit of detection (LOD) was defined as the lowest input DNA concentration at which at least one out of three replicates tested positive.

The SINE marker detected significantly higher eDNA concentrations than the 16S marker in a field survey conducted in *A. japonica* habitats. The average copy number per liter of water was approximately 170 times higher for the SINE marker (mean = 17,458.6 copies/L) than for the 16S marker (mean = 102.2 copies/L) (LMM, *p* = 1.06 × 10□¹¹; Fig. 3, Fig. S1). The eDNA concentrations of the two markers were strongly positively correlated (Model II regression, *R* = 0.931, *p* = 1.78 × 10□¹²; Fig. S2).

**Figure 3.**
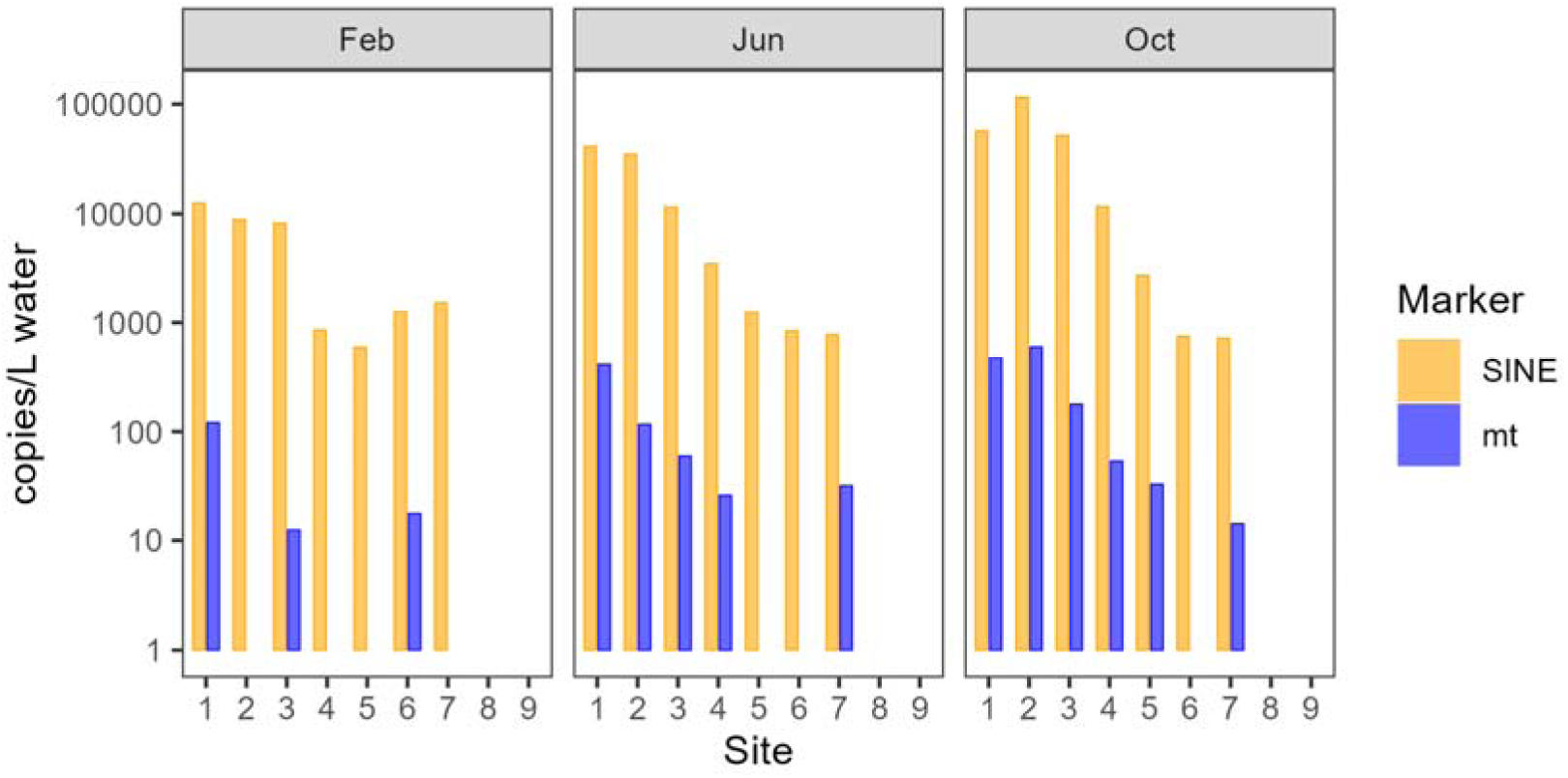
Mean copy numbers per liter of water for the SINE (UnaSINE1) and mitochondrial 16S (mt) marker at each sampling point in the Sumiyoshi River.

Across all survey periods, the eDNA of *A. japonica* was detected using both markers at least once at seven of the nine sites, excluding the two most upstream sites. The mitochondrial 16S marker was detected at 14 points (32 bottle samples), whereas the SINE marker was detected at 21 points (62 bottle samples). The detection probability per survey point was significantly higher for the SINE marker (*p* = 0.023, McNemar’s test; Fig. 4). At sites where mitochondrial marker were detected, SINE marker were significantly more likely to be positive in all biological replicates (*p* = 6.151 × 10□□, McNemar’s test). The CV values for both technical and biological replicates were significantly lower for the SINE marker than for the mitochondrial marker (Wilcoxon rank sum test, technical replicates: *p* = 3.21 × 10□□, biological replicates: *p* = 0.00812; Fig. S3).

**Figure 4.**
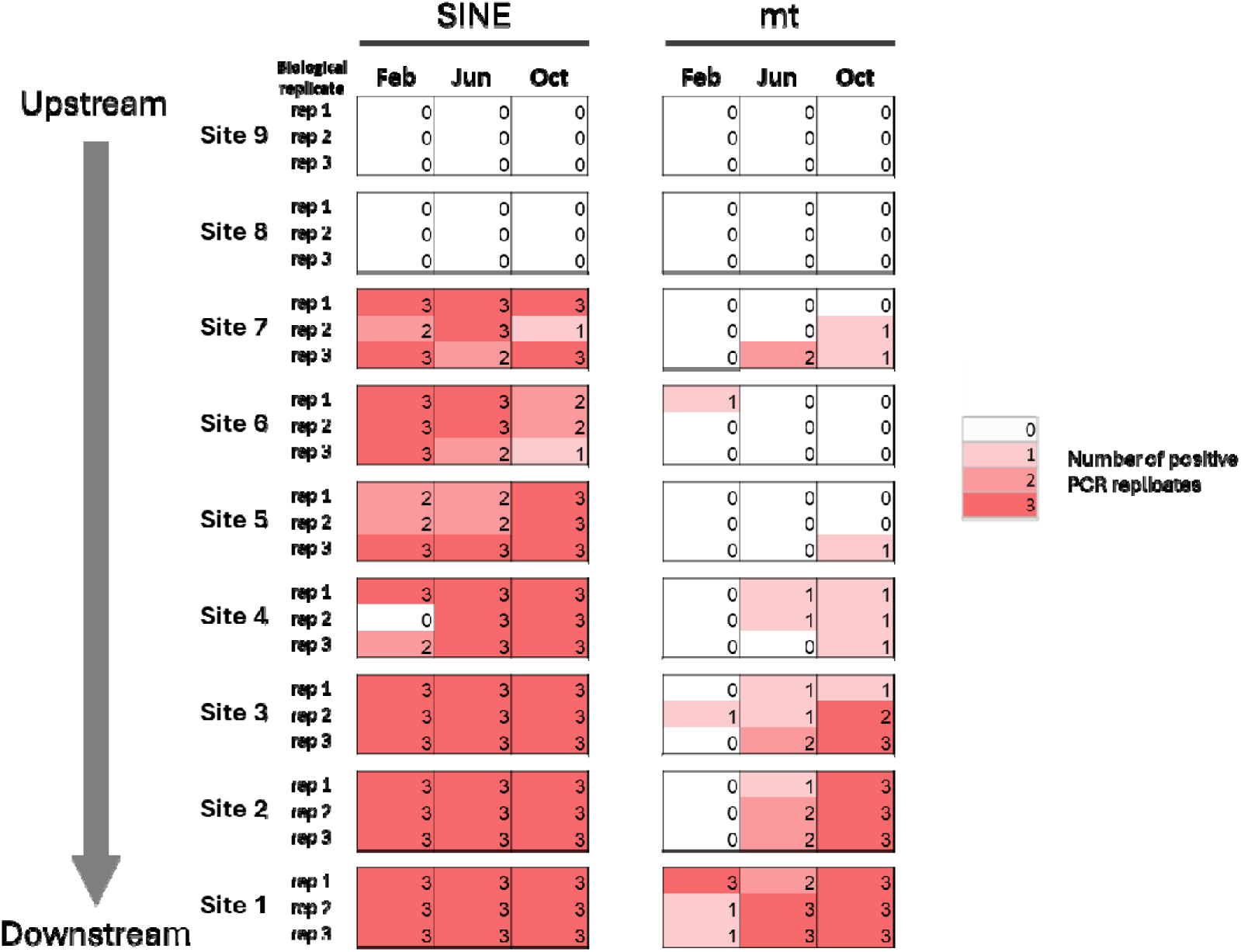
Number of PCR-positive reactions from each bottle sample collected at the Sumiyoshi River. Three bottle samples (biological replicates) were collected at nine sites (Sites 1–9) for each sampling period (February, June, and October). Filled cells indicate bottle samples in which at least one out of three PCR replicates (technical replicates) tested positive.

Significant differences in eDNA concentrations between sites were observed for both markers (LM, mitochondrial marker: *p* = 0.0476, SINE marker: *p* = 4.33 × 10□□). However, no significant inter-site differences were detected using the mitochondrial 16S marker in multiple comparisons with Tukey’s adjustments. In contrast, the SINE marker identified three distinct eDNA concentration groups: a high-concentration downstream group (Sites 1 and 2), an intermediate group (Sites 3 and 4), and a low-concentration upstream group (Sites 5, 6, and 7) (Fig. S4). Sites 3 and 4 overlapped with the downstream and upstream groups, respectively. Temporal variations in eDNA concentrations were also significant for both markers (LMs: mitochondrial marker, *p* = 0.0441; SINE marker, *p* = 0.0203). The eDNA concentrations in February and October were significantly different, whereas that in June showed no significant difference compared with that in the other months.

## 4. Discussion

We successfully developed an eDNA assay targeting the *Anguilla*-specific SINE family, UnaSINE1, and demonstrated that it provides higher sensitivity and accuracy than the commonly used assay utilizing mitochondrial markers. To the best of our knowledge, this is the first study to detect retrotransposons in water samples. The conservation of *Anguilla* species is an urgent issue, and accurate monitoring of their distribution and biomass in natural habitats is crucial for effective conservation.

In the sensitivity test using *A. japonica* gDNA, the LOD and LOQ of the SINE marker were both two orders of magnitude lower than those of the mitochondrial marker, suggesting that the copy number of *Anguilla*-specific UnaSINE1 per cell was at least 100 times greater than that of mitochondrial genes.

The presence of eel species was confirmed during field surveys conducted in a freshwater river where *A. japonica* inhabits, with both markers at all sites except the two most upstream sites. In contrast, there is a possibility that a weir more than 3 m high, built upstream of Site 7, hinders eel migration and restricts their habitat. The strong positive correlation between the eDNA concentrations of the two markers suggested that the eDNA concentration of each marker reflects the biomass of *A. japonica* in its habitat. However, the SINE marker yielded approximately 170 times more copies per liter of water than the mitochondrial marker, indicating a substantial difference in environmental eDNA abundance due to the higher copy number of the SINE marker in the cells.

The use of the SINE marker significantly improved the detection sensitivity. While there were no bottle samples in which only the mitochondrial marker was detected, 30 bottle samples tested positive only for the SINE marker. Additional detection of *A. japonica* eDNA was performed at four sites in February, two in June, and one in October. These findings suggested that false negatives for the mitochondrial marker, likely caused by low individual density or low eDNA release rates, were resolved using the SINE marker. Furthermore, the eDNA concentrations at sites where new detections were made using the SINE marker exceeded those at the upstream sites. This suggested that these detections were not false positives caused by downstream transport of eDNA from upstream populations. Thus, using the SINE marker is particularly effective for detecting low-density populations in field surveys (Kume et al., 2021). Given that *Anguilla* species undergo long-distance spawning migration, surveys in oceans, where eel densities are much lower than those in rivers, are essential (Bonhommeau et al., 2010; Noda et al., 2021). The high sensitivity of the SINE marker assay is expected to be particularly beneficial in such investigations.

False negatives in eDNA analysis are primarily caused by the failure to capture molecular markers during sampling or subsequent analytical processes. In this study, the biological and technical replicates were not always consistent, even at the sites where the mitochondrial marker was detected, with some replicates showing no detection. This is likely due to the insufficient eDNA concentration of the mitochondrial marker. However, all biological replicates tested positive at sites where the mitochondrial marker was positive when using the SINE marker. Furthermore, only one discrepancy among the biological replicates was observed throughout the study period (February, Site 4). This suggested that abundant molecular markers in the water samples reduced the likelihood of false stochastic negatives during sampling. The detection error rate in technical replicates was also significantly lower for the SINE marker than that for the mitochondrial marker, indicating improved assay precision. The CV values were significantly lower for both biological and technical replicates with the SINE marker, suggesting that this marker enhances the reliability of quantitative eDNA analysis. This improvement enables more accurate presence/absence determinations and more robust quantitative comparisons of eDNA concentrations.

In the present study, multiple comparison tests using the SINE marker revealed three significantly different eDNA concentration groups: higher concentrations downstream, lower concentrations upstream, and intermediate concentrations in between. Lower eDNA concentrations in the upstream regions suggested a negative correlation between *A. japonica* population density and distance from the river mouth, reflecting the species’ habitat preference (Itakura et al., 2020b). A series of river-crossing structures, which may have hindered the upstream migration of *A. japonica*, were present between Sites 3 and 4. In contrast, the mitochondrial marker did not detect a clear spatial gradient in eDNA concentration. This is likely due to the higher variability and lower detection probability associated with the mitochondrial marker. The lower eDNA concentrations observed in February for both markers suggested that the reduced activity of *A. japonica* during winter, driven by low water temperatures, resulted in a lower eDNA release (Itakura et al., 2015). Only three sites yielded positive detections with the mitochondrial marker in February, whereas the SINE marker detected eDNA at all sites, except for the two most upstream locations. This suggested that the SINE marker can mitigate false negatives in seasonal surveys, particularly when the activity of the target species decreases. In summary, the improvement in sensitivity revealed that the distribution of eels was clearly restricted upstream of Site 8, whereas the improvement in accuracy demonstrated a significant decrease in eel biomass upstream of Site 3. Artificial weirs might have been the cause in both cases. Because eels possess a high climbing ability, the impact of small-scale river-crossing structures on their upstream migration has not received much attention (Matsushige et al., 2021). However, based on the results of this study, it is necessary to reconsider this perception.

Improving sensitivity and accuracy achieved by changing the assay target is advantageous because it does not increase the workload or reagent costs associated with eDNA collection and PCR analysis. Sufficient markers can be recovered with a smaller filtration volume during sampling, which is expected to reduce processing time and minimize the impact of inhibitory substances present in environmental water. Reducing the template volume per reaction and the number of PCR replicates may help conserve samples and reagents.

One of the challenges of using SINEs as markers for eDNA analysis is that their applicability is currently limited to specific organisms. Mitochondrial and ribosomal genes exist in all organisms and can generate paralogues through mutations. In contrast, SINE families arose through rapid copy number expansion during evolution. Consequently, not all species or taxonomic groups possess unique SINE families (Frengen et al., 1991; Takasaki et al., 1996). Additionally, when a SINE family originates before species divergence, it may be shared among closely related species with a common ancestor, thereby lacking species specificity. The classical method for isolating retrotransposons involves screening genomic libraries using RNA probes, which require more effort than mitochondrial markers, where sequence information can be easily obtained through PCR amplification and Sanger sequencing. These factors present challenges to the development of eDNA markers.

For UnaSINE1, *in silico* PCR confirmed its amplification in multiple *Anguilla* species, suggesting that the SINE family originated from a common ancestor. Although species-specific UnaSINE1 variants have been identified, designing primers and probes to maximize the number of detectable sequences requires a balance between specificity and sensitivity. Consequently, we developed an assay targeting *Anguilla* at the genus level rather than using species-specific markers. This approach limits species-level identification in regions where multiple *Anguilla* species coexist. However, it remains highly effective in areas where only one species is present, such as our study sites.

Given that *A. marmorata* gDNA yielded Ct values similar to *A. japonica*, this assay could likely be performed for other *Anguilla* species. Future studies using whole-genome sequencing and repeat tome analyses may lead to identifying species-specific SINE families. As genomic databases expand beyond model organisms, using nuclear repeat elements, such as SINEs, as eDNA markers is expected to become increasingly viable.

## Acknowledgements

We are grateful to Dr. Nobuko Ohmido for helpful comments. This study was funded by a Japanese Society for the Promotion of Sciences (JSPS) Grant-in-Aid for Fellows to ITH (Grant No. JP23KJ1569). We used DeepL Translate (https://www.deepl.com/ja/translator) for translation and grammatical checks. We would like to thank Editage (www.editage.jp) for English language editing.

## Data Availability Statement

The data that supports the findings of this study are detailed in the article’s figures and tables and are available in the Supporting Information of this article.

