## Supplemental Information for "Significant improvement of environmental DNA assay by targeting retrotransposon sequences characteristic to *Anguilla* eels"

**Table S1.** Standard sequence used in the UnaSINE1 assay.

Sequence (5ʹ-3ʹ)

GGAGTGTAGCACAGTGGGTAAGGAACTGGGCTTGTAACCGAAAGGTCGCAGGTTCGATTCCCAGGTAAGGACACTGCCGTTGTACCCTTGAGCAAGGTACTCAACCGGAATTGATTCAGTATATATCCAGCTGTATAAATGGATACAATGTAAAATGCAATGTAAAAGTTGTGTAAGTCGCT

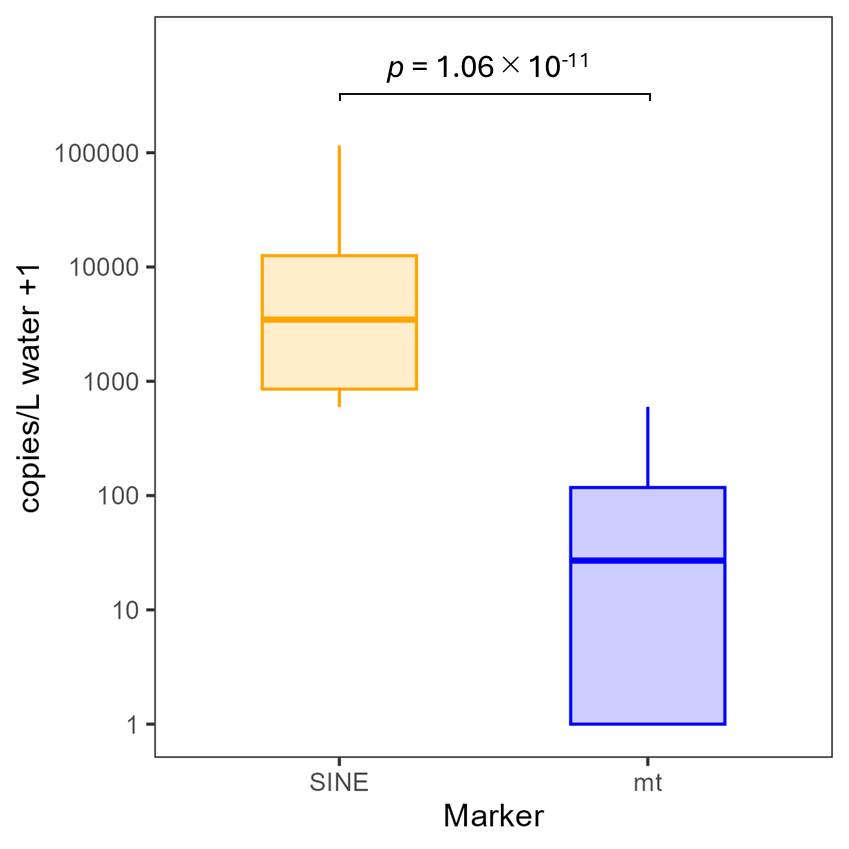

**Figure S1.** Differences in copy numbers of the UnaSINE1 (SINE) and mitochondrial 16S (mt) markers per liter of water in the Sumiyoshi River. eDNA marker concentrations per liter were log10-transformed (+1) and compared using a linear mixed model. The explanatory variable was the marker type, and random effects included the sampling site and month.

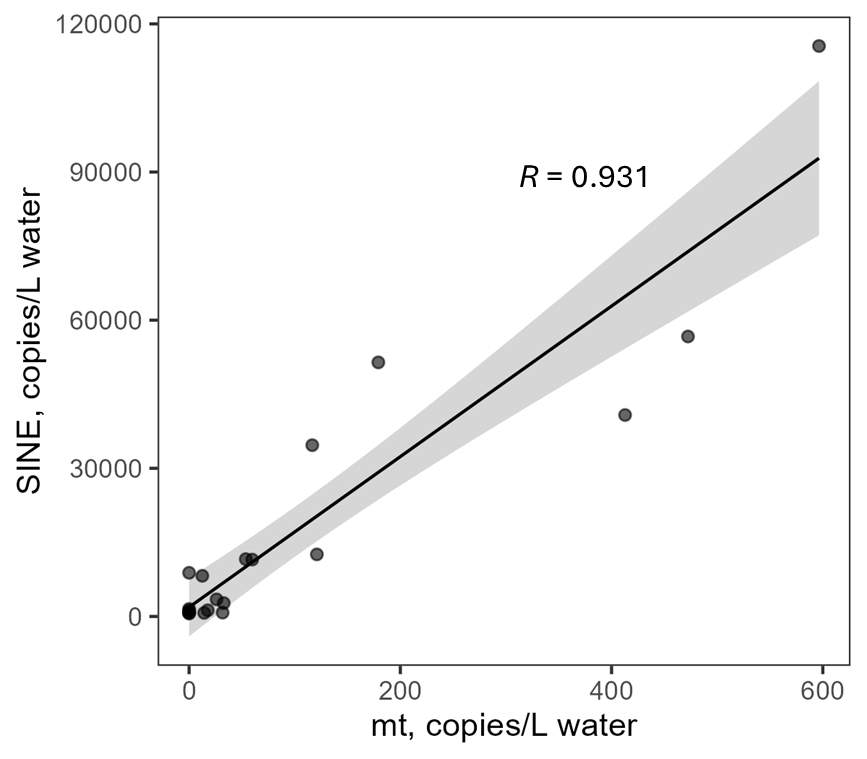

**Figure S2.** Scatter plot of mean copy numbers per liter of water for the UnaSINE1 (SINE) and mitochondrial 16S (mt) markers (3 months × 7 sites = 21 points) in the Sumiyoshi River. Sites 8 and 9, where eels were never detected during the survey period, were excluded from the analysis. The black line represents the regression line fitted with Model II regression, and the gray shading indicates the 95% confidence interval.

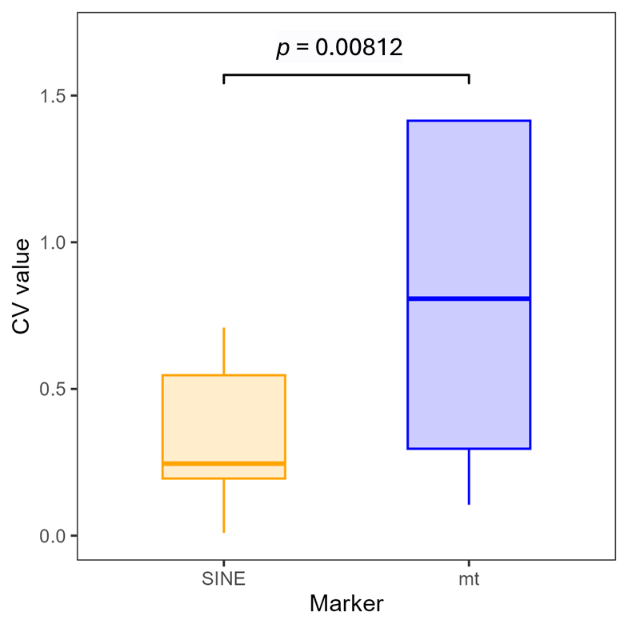

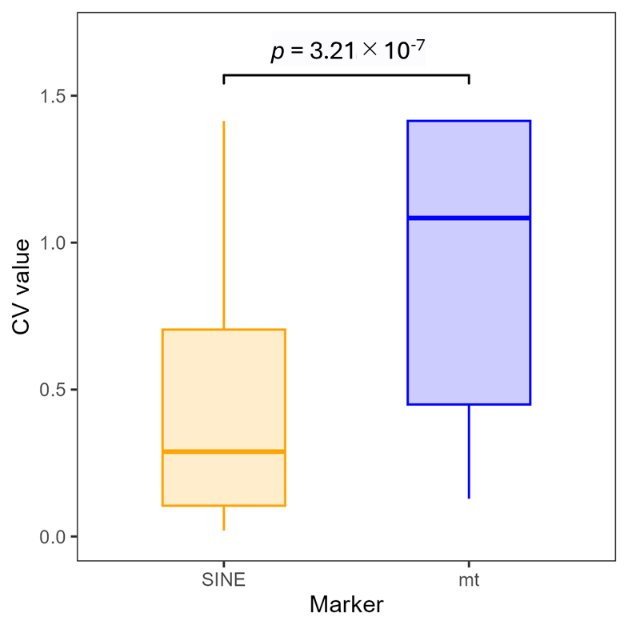

**Figure S3.** Comparison of coefficient of variation (CV) values for eDNA concentrations in biological replicates (bottle samples, left) and technical replicates (PCR reaction, right). The Wilcoxon signed-rank test was used to evaluate significant differences between UnaSINE1 (SINE) and mitochondrial 16S (mt) markers.

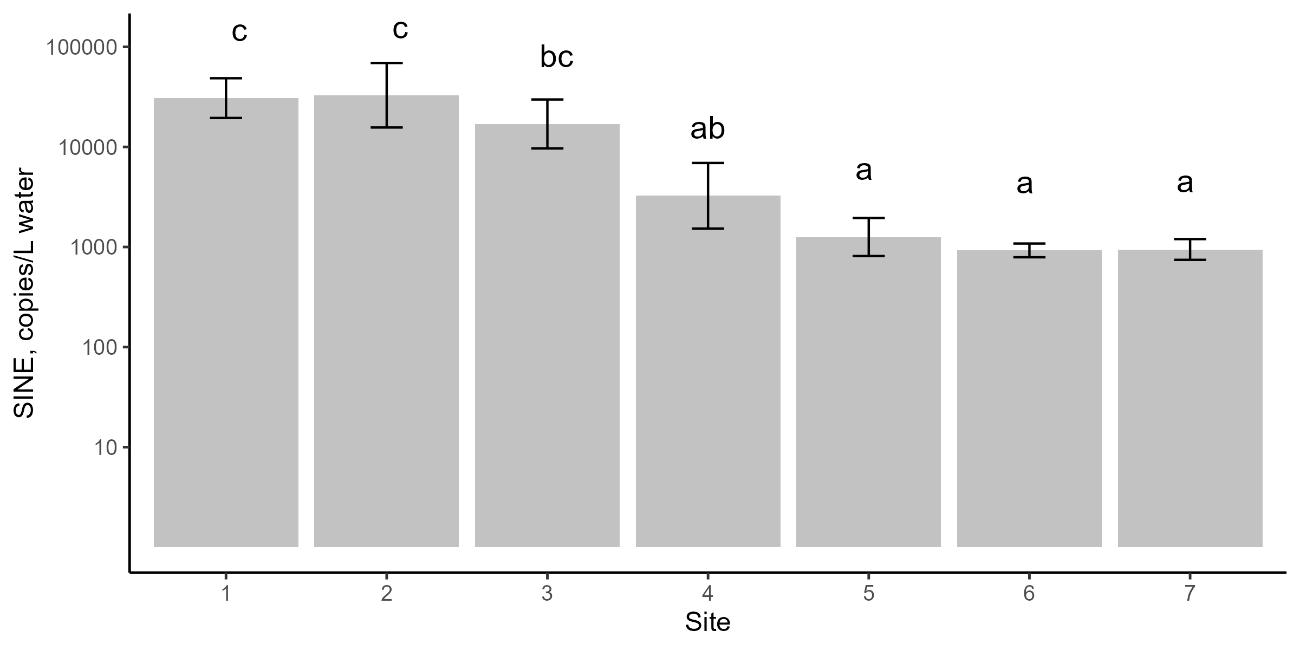

**Figure S4.** Mean copy numbers per liter of water for the UnaSINE1 (SINE) marker at Sites 1–7 in the Sumiyoshi River. Sites with different letters indicate significant differences based on Tukey-adjusted multiple comparisons (*p* < 0.05).
